# Neural effects of continuous theta-burst stimulation in macaque parietal neurons

**DOI:** 10.1101/2020.12.07.414482

**Authors:** Maria C. Romero, L. Merken, P. Janssen, M. Davare

## Abstract

Theta-burst transcranial magnetic stimulation (TBS) has become a standard non-invasive technique to induce offline changes in cortical excitability in human volunteers. Yet, TBS suffers from a high variability across subjects. A better knowledge about how TBS affects neural activity in vivo could uncover its mechanisms of action and ultimately allow its mainstream use in basic science and clinical applications. To address this issue, we applied continuous TBS (cTBS, 300 pulses) in awake behaving rhesus monkeys and quantified its after-effects on neuronal recordings and behavior. Guided by anatomical MRI, we recorded single-cell activity in parietal area PFG during passive fixation of real-world objects. Overall, we observed a pronounced, long-lasting and highly reproducible reduction in neuronal excitability after cTBS in individual parietal neurons, with some neurons exhibiting periods of hyperexcitability during the recovery phase. We applied the same stimulation protocol during visually-guided grasping of objects, and observed a significant grasping impairment. These results provide the first experimental evidence on the effects of cTBS on single neurons in awake behaving monkeys.

## INTRODUCTION

Repetitive Transcranial Magnetic Stimulation (rTMS) protocols, such as continuous Theta-Burst stimulation (cTBS), represent a non-invasive way to reduce cortical excitability in human volunteers (Huang et al., 2005) and to explore neuroplasticity in several patient populations (Edwards et al., 2006; Oberman et al., 2010; Orth et al., 2010; Suppa et al., 2011, 2014a,b; Koch et al., 2012; Munneke et al., 2013; Opie et al., 2013; Chuang et al. 2014; Mori et al., 2014). However, a number of major hurdles prevent its widespread application in basic research and in clinical care. For example, despite more than 10 years of studies in human volunteers, it is still unclear how neural activity changes in the cortex after cTBS (Pitcher et al., 2020). It is generally assumed that the reduction in neuronal excitability may resemble the changes observed in long-term depression (LTD; Fitzgerald et al., 2006; Thut and Pascual Leone, 2010; Vlachos et al., 2012), but no study has ever tested in vivo whether this is true. The reduction in the amplitude of the motor evoked potential (MEP) after cTBS also grows over time (Huang et al., 2005), which does not resemble the immediate reduction in neuronal excitability after conditioning observed in LTD. Moreover and more importantly, the physiological effects of cTBS are notoriously variable, with inhibitory effects in some subjects and facilitatory effects in other subjects (Hamada et al., 2013). Several factors could contribute to this variability (reviewed in Suppa et al., 2016). Genetic variation, differences in the intracortical network activated by the TMS pulses (measured in the so-called late I-waves), previous levels of activity, circadian effects and differences in the brain state may all contribute to the inter- and intra-individual variability reported in cTBS studies.

We sought answers to the aforementioned questions by recording single-neuron activity before and after cTBS in parietal cortex of awake behaving rhesus monkeys. We probed the neuronal excitability using single-pulse TMS, and measured how this excitability changed up to two hours after cTBS. To the extent possible, we standardized the factors potentially contributing to the known variability associated with cTBS: the positioning of the coil on the skull, the level of ongoing motor activity, the stimulation hour and the brain state were equalized as much as possible across sessions and animals. We observed highly reproducible changes in neuronal excitability, in which individual neurons would progress through phases of hypo- and hyperexcitability, in some cases followed by recovery. At the population level, the reduction in neuronal excitability grew over time, reaching its maximum 30-40 min after cTBS, consistent with studies in humans. Behaviorally, we measured a significant deficit in object grasping, the dynamics of which mirrored the changes in excitability detected in the neurons. Thus, in a standardized experimental setting, cTBS induces reproducible and long-lasting changes in neuronal excitability and behavior.

## TECHNIQUES AND METHODS

### Subjects and surgical procedures

Four adult male rhesus monkeys (Macaca mulatta; monkey Y, 12 kg; monkey A, 8 kg; monkey P, 10 kg; monkey D, 7.5 kg) were trained to sit in a primate chair. A head post (Crist Instruments) was then implanted on the skull with ceramic screws and dental acrylic. For this and all other surgical protocols, monkeys were kept under propofol anesthesia (10mg/kg/h) and strict aseptic conditions. All experimental procedures were performed in accordance with the NIH’s Guide for the Care and Use of Laboratory Animals and the EU Directive 2010/63/EU, and approved by the Ethical Committee at KU Leuven. Intensive training in passive fixation and visually-guided grasping (VGG) began after 6 weeks of recovery. Once the monkeys had achieved an adequate level of performance, a craniotomy was made in monkeys A and Y, guided by anatomical Magnetic Resonance Imaging (MRI), over area PFG of the right hemisphere. An exhaustive description of this protocol has been detailed elsewhere (Romero, Davare et al., 2019). The recording chamber was implanted at a 45 deg angle with respect to the vertical, allowing oblique penetrations into the parietal convexity (Figure 1A). To confirm the recording positions, glass capillaries were filled with a 2% copper sulfate solution and inserted into a recording grid at five different locations during structural MRI (slice thickness: 0.6 mm). In all four animals, two guiding rods were precisely fixed to the skull with dental acrylic based on the calculated stereotactic coordinates, allowing a highly reproducible positioning of the TMS coil across sessions. With these rods in place, the coil was kept at an angle of approximately 90 deg with respect to the recording chamber in monkeys Y and A, inducing a posterior-anterior (PA) current over PFG. In monkeys P and D, the coil was centered on area PFG, mimicking the coil angle and orientation used in the electrophysiology experiments. To estimate the center of stimulation, in monkeys Y, A and P, we used anatomical MRI and Computed Tomography (CT scan) to build 3D printed models of the skull and implant (Figure 1A). Based on the MR-CT co-registered images, we calculated that the TMS coil was placed approximately at a distance of 15 mm from (above) the parietal convexity. In monkey D, we estimated the center of stimulation based on anatomical MRI with a dummy coil placed over the guiding rods, also targeting the parietal convexity. As for the other three animals, the distance between the coil and the cortex was 15 mm.

**Figure 1.**
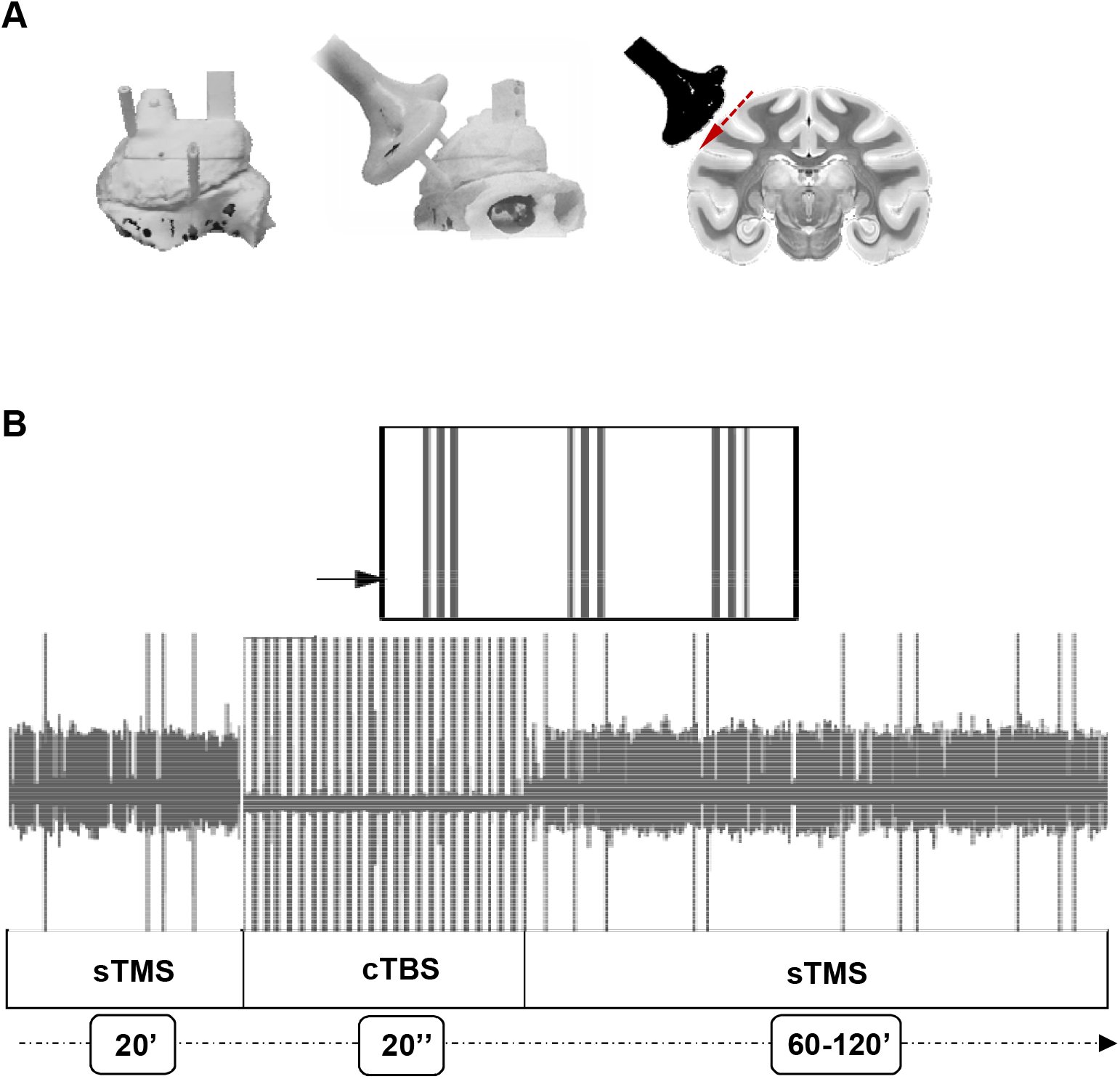
Brain targeting and stimulation protocol. **A.** Three-dimensional models of the monkey’s skull (left: top view, right: lateral view) and coronal view of the monkey brain indicating the location of the TMS coil with respect to parietal area PFG. The red dashed line indicates the trajectory of the electrode during recordings. For every experiment, a 25 mm figure-of-eight TMS coil (in black) was rigidly anchored to the monkey’s skull to allow a precise and reproducible coil positioning across experimental sessions. Two guiding rods were attached to the monkey’s head implant based on MRI estimations of the cortical target coordinates. **B.** Example of the raw signal recorded during a typical stimulation session in Experiment 1. The high voltage, saturated peaks indicate the stimulation time stamps. TMS was administered in three different epochs corresponding, in this order, to: single-pulse TMS (sTMS) applied at light onset (20 min), cTBS (20 sec), and again, sTMS (60-120 min).

### Stimulation protocol

For this study, we combined two different TMS protocols: single-pulse TMS (sTMS) and continuous Theta-Burst Stimulation (cTBS). In the first experiment, we performed extracellular recordings before and after cTBS to investigate the changes in neuronal excitability, assessed with sTMS, during passive fixation. In the second experiment, we studied the effect of cTBS on motor behavior by measuring its effect on grasping times (GT) during visually-guided object grasping (VGG).

### Experiment 1. cTBS effect on individual neurons: electrophysiological recordings

Two male rhesus monkeys were used for this experiment (monkeys Y and A). To optimize the stability of the recordings (total duration 1 to 3 hours per neuron), the animals performed a passive fixation task while sitting up right with the head fixed. With this setup, a single object (large sphere; diameter: 35 mm) was placed on a frontal plate, located 30 cm away from the animal. In the beginning of the trial, the monkeys remained in total darkness for a variable time period (intertrial interval: 2000-3000). Next, a red laser was projected at the base of the object, indicating the fixation point. If the animal maintained fixation inside an electronically-defined window (± 2.5 deg around the fixation point) for 500 ms, the object was illuminated (light onset) until the end of the trial (1300 ms). Whenever the monkey maintained a stable eye fixation, it received a drop of juice as reward.

Prior to the experiment, we determined the resting motor threshold (rMT) for each animal as the lowest stimulus intensity at which TMS over primary motor cortex (M1) produced a contralateral finger twitch while the monkey held its hand in the resting position. For this measurement, the TMS coil was handheld over M1, at a distance of approximately 15 mm from the surface of the brain, reproducing the coil distance adopted during the experiments.

In each recording session, the coil was placed over the guiding rods, tangential to the skull, staying firmly anchored to the chair by means of an adjustable metal arm. First, we applied sTMS at 120% of the rMT (Figure 1B), aligned to light onset and randomly interleaved with no-stimulation trials, while advancing the electrode and searching for well-isolated units. We used a Magstim Rapid Stimulator (Magstim, UK), which applied biphasic pulses (100 μs rise time, 250 μs duration; 120% rMT = 70% of maximum stimulator output) by means of a 55 mm figure-of-eight coil. As for previous experiments in our group, we employed this protocol to localize the center of stimulation, which was determined as the region under the center of the coil showing a significant sTMS-evoked effect in single neurons. With our technique, we isolated individual PFG neurons and stabilized the recordings to guarantee a reliable long-lasting monitoring of their activity. In each recording session, we first waited a variable amount of time (40-60 min) to acquire the highest possible level of stability in the recorded signal and then measured the baseline neuronal activity and the sTMS-evoked response for 20 min. In order to prevent a possible overheating of the coil during this period, a gel-filled cool pack was placed around the handle, embracing the figure-of-eight-shaped head of the coil (Nexcare; 3M Company, Minnesota, USA). Next, we applied cTBS (Figure 1B) following the protocol described by Huang and colleagues (Huang et al., 205). In our cTBS paradigm, 50 Hz triplets were delivered at 200 ms intervals (300 pulses in total) for 20 sec at 80% of the rMT (47% of the maximal intensity). To prevent coil overheating during this phase, the gel-filled cool pack was now substituted by two round cool bags containing dry ice and placed over the loop ends of the coil. Phase three started immediately after cTBS, and consisted of a second period of combined sTMS and electrophysiological recordings, at 120% of the rMT for up to two hours post-cTBS to measure changes in neuronal excitability. For the entire duration of the experiments, none of the animals showed any noticeable side-effect to cTBS, performing their task without signs of distress.

We recorded single-unit activity in PFG using tungsten microelectrodes (impedance: 1 MΩ at 1 kHz; FHC) inserted through the dura by means of a 23-gauge stainless steel guide tube and a hydraulic microdrive (FHC, USA). Following the artifact reduction strategy proposed by Mueller and colleagues (Mueller et al., 2014), we used diodes and serial low-gain amplification to clip the artifact generated by the magnetic pulses, which prevented amplifier saturation. For this, we modified a regular BAK Electronics preamplifier (Model A-1; BAK Electronics, USA) by connecting two leakage diodes (BAS45A) anti-parallel between the signal lines and ground before each stage of amplification. The initial front-end of the headstage remained unmodified to maintain the high-input impedance. With these settings, the duration of the evoked TMS artifact in our signal ranged from 8 to 12 ms (Romero, Davare et al., 2019). Neural activity was amplified and filtered (300-5,000 Hz) following a standard recording protocol for spike detection. Using a dual time-window discriminator (LabVIEW and custom-built software), we isolated individual neurons and the TMS artifact, which was detected online and subtracted from the neural data. In addition, we recorded the entire raw signal (after filtering) for further analyses. Finally, we monitored the right eye position using an infrared-based camera system (Eye Link II, SR Research, Canada) sampling the pupil position at 500 Hz.

### Experiment 2. cTBS effect on behavior: visually-guided grasping task

Experiment 2 was designed to assess the effect of cTBS on motor behavior. Therefore, no electrophysiological recordings were performed during these sessions.

Two male rhesus monkeys (Macaca mulatta; monkeys P and D) were trained for this experiment. All surgical and chair training protocols were identical to those described for experiment 1. However, in these animals, the behavioral training consisted of a motor task (VGG task). In the VGG task, for each recording session, the hand ipsilateral to the recording chamber remained restrained within the chair, while the contralateral hand was placed on a resting device. A single grasping object (a cylinder with a diameter of 35 mm and a height of 38 mm) was located in front of the monkey, at the same distance as the display in the passive fixation task (30 cm away from the animal). The resting position of the hand, the start of the reach to grasp movement and the pull of the object were detected by fiber-optic cables. During the task, the monkey had to place the hand contralateral to the recorded hemisphere in the resting position in complete darkness to initiate the sequence. After a variable time (intertrial interval: 2000-3000 ms), a red laser projected a fixation point at the base of the object. If the animal maintained its gaze inside the electronically-defined fixation window (+/− 2.5 deg) for 500 ms, the object was illuminated. Following a variable delay (900-1100 ms), a visual GO cue (dimming of the laser) instructed the monkey to lift the hand from the resting position, and reach, grasp, lift and hold the cylinder for a variable interval (holding time, 500-900 ms). Whenever the monkey performed the whole sequence correctly, it received a drop of juice as reward. For every session, we measured the time elapsed between the start of the movement and the lift of the object (grasping time: GT).

Prior to the experiments, we conducted a pilot study (4 sessions per object) to quantify the baseline performance of each monkey. During this period, no cTBS was applied. Next, the experimental phase began, in which cTBS was delivered using a Magstim Rapid Stimulator (Magstim, UK) and a custom-built figure-of-eight coil for animal use (D25 mm; 55 mm external diameter; same as for experiment 1). As for experiment 1, the cTBS paradigm consisted of 50 Hz triplets repeated at 200 ms intervals (300 pulses) for a total time of 20s. During the study, we recorded the animal’s performance in three different stimulation conditions: no-stimulation, low-intensity stimulation (40% of the rMT) and high-intensity stimulation (80% of the rMT). The three conditions were randomly interleaved across sessions. In a typical stimulation session, the monkey first performed the VGG task for 20 min (baseline acquisition). Next, low- (40% of the rMT) or high-intensity (80% of the rMT) cTBS was applied for 20 seconds. During this period, the behavioral response of the monkey was ignored and the trials performed (if any), discarded from the final analyses. After the application of cTBS, the behavioral task resumed, monitoring the effect of stimulation for up to 120 min. To guarantee reliable results, a minimum of 2000 trials per condition were collected across sessions (minimum of 3 sessions per stimulation level). As for experiment 1, both monkeys in this study showed no side-effects after cTBS stimulation and performed the grasping task without any signs of distress.

### Data analysis

All data analyses were performed in MATLAB (MathWorks, Massachusetts, USA; code availability: DRYAD database). For the high- and low stimulation trials of experiment 1, the neural activity was aligned on the sTMS pulse delivered at light onset. Also, for comparison, the no-stimulation trials were aligned on the same time bin. Net neural responses were then calculated as the average firing rate recorded after sTMS minus the baseline (spike rate calculated from the mean activity of the cell in the 800 ms interval preceding TMS).

We created line plots comparing the average response (spikes/sec) of every cell during no-stimulation and stimulation(sTMS) trials recorded in the pre-cTBS epoch. The same analyses were then repeated in 10 min epochs post-cTBS, in order to assess in detail how the response of PFG neurons to sTMS – our measure of neuronal excitability – changed after cTBS. Since we expected that cTBS would induce cortical inhibition, we searched for PFG neurons showing excitatory responses to sTMS.

To determine the significance of the sTMS-evoked effect on individual neurons, we compared the cell responses observed in the first 40 ms after light onset in the stimulation condition to those in the no-stimulation condition (two-sided Wilxocoxon ranksum test). To identify neurons with task-related activity, we ran a two-sided Wilcoxon ranksum test comparing the pre- and post-light onset responses in two different intervals (early task activity: 20-100 ms post-stimulus; later task activity: 120-520 ms post-stimulus) in the no-stimulation condition. Finally, to assess the effect of cTBS on neuronal excitability, we compared the average net sTMS-evoked response in every 10-min epoch after cTBS with the same response pre-cTBS (baseline sTMS net response, two-sided Wilcoxon ranksum test). For all neurons recorded for more than one hour, we compared the raw traces to verify that the neuron was not lost during the recordings.

For experiment 2, we measured the effect of cTBS on motor performance by analyzing the grasping time (GT, i.e. the time elapsed between the lift of the hand and the pull of the object) in all correct trials of the VGG task. Because of the large intersession variability in grasping behavior observed for both monkeys, GTs were normalized according to the following equation:

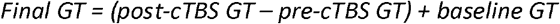

where the pre-cTBS GT corresponds to the average GT obtained before applying cTBS and the baseline GT refers to the average GT measured during the four sessions corresponding to the pilot study (without cTBS stimulation).

To evaluate the effect of cTBS across time, the average GT was calculated in 10 min intervals, which also allowed direct comparisons between the observed behavioral effect in the VGG task and the cTBS-induced effect on individual neurons, computed in Experiment 1.

To determine the cTBS effect on behavior, we calculated two-sided Wilcoxon signed-rank tests (corrected for multiple comparisons) on high- and low stimulation trials compared to no-stimulation in each 10 min epoch. Similarly, we performed a one-way ANOVA on high-, low- and no-stimulation trials to determine the effect of stimulation intensity on GT. Finally, to quantify the effect of cTBS on GT over time, we ran a repeated-measures ANOVA with time and stimulation as factors, corrected for multiple comparisons (Tukey’s test).

## RESULTS

We recorded the activity of 86 single neurons in parietal area PFG of two monkeys (51 neurons in monkey Y; 35 in monkey A) before and after cTBS during passive fixation. In two other monkeys (P and D, one of which was used in the Romero, Davare et al., 2019 study), we measured the behavioral effect of cTBS applied over parietal cortex using an object grasping task.

### Effect of cTBS on individual neurons

Before applying cTBS, we tested the excitability of each neuron using single-pulse TMS (sTMS) administered at light onset above the object. Figure 2 illustrates, with two examples, the typical results in this study (Experiment 1). The first example neuron (Figure 2A) generated a brief burst of action potentials almost immediately after sTMS (top row), but did not respond to light onset in the absence of sTMS. After 20 sec of cTBS, however, the excitability of this neuron was markedly reduced (second row, sTMS-evoked response pre-compared to post-cTBS, p = 1.062 e-04, Wilcoxon test). Indeed, in the first 10 min post-CTBS, the activity in high-stimulation trials did not differ anymore from the activity in no-stimulation trials (p = 0.070). Over the subsequent time intervals, the excitability of this neuron gradually recovered (no-stimulation compared to stimulation trials, p = 6.011e-07, p = 4.567e-11, p = 1.353e-07, p = 2.173e-06 for the 20, 30, 40 and 50 min interval, respectively), but only after 60 min did the excitability of the neuron return to the pre-cTBS level (sTMS-evoked response pre-compared to 60 min post-cTBS, p = 0.103). Thus, cTBS caused a marked and immediate reduction in excitability in this parietal example neuron, which recovered over the course of one hour.

**Figure 2.**
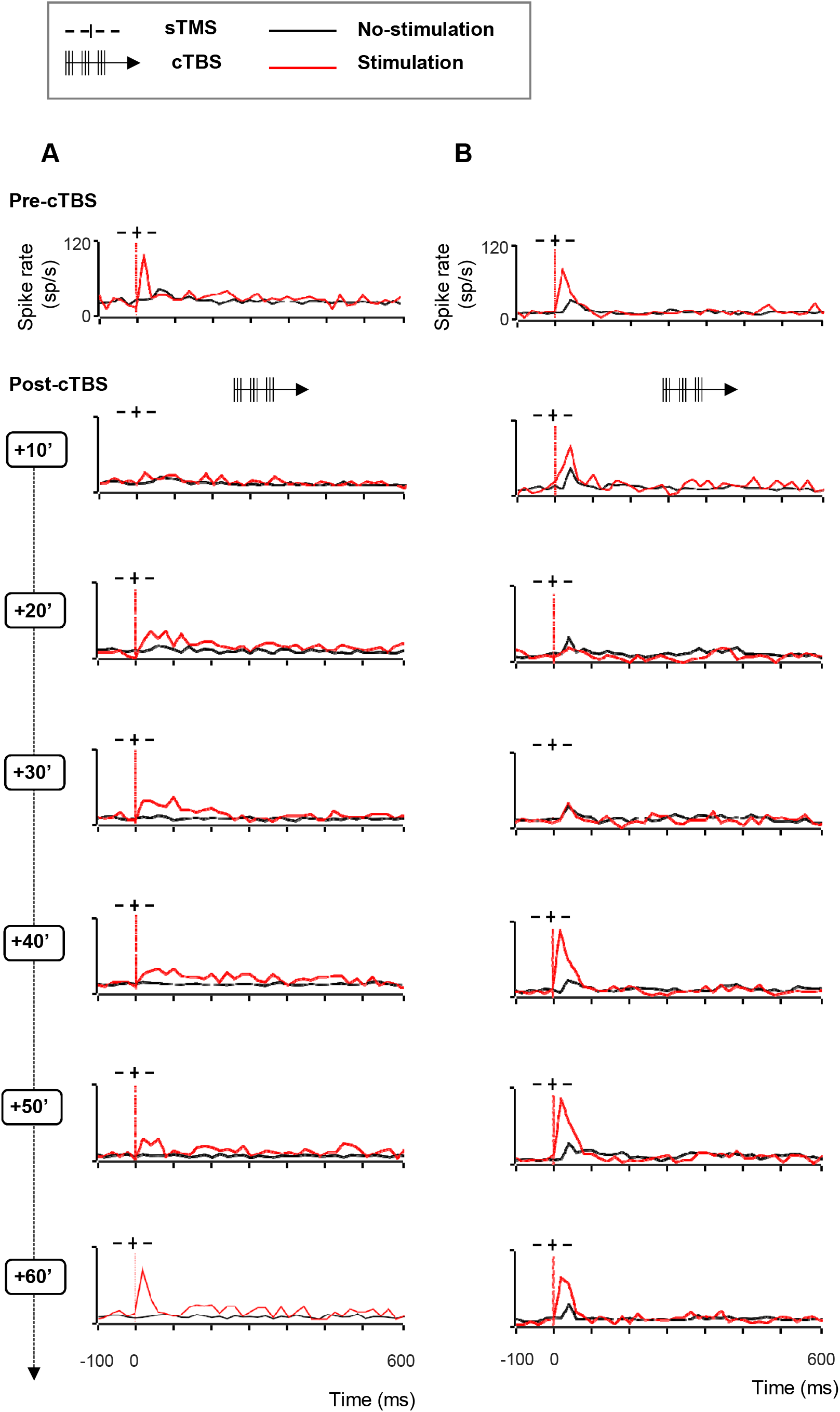
Effect of cTBS on neuronal excitability, example neurons with recovery. We tested the excitability of each neuron using single-pulse TMS (sTMS) administered before and after cTBS. Stimulation (red line plots) and no-stimulation trials (black line plots) were randomly interleaved during the passive fixation task. The red, dotted line indicates the sTMS onset (aligned to light onset). **A.** Spike rate of an example neuron exhibiting a short-lasting, excitatory response to sTMS (top row). cTBS caused a marked and immediate reduction in excitability (row 2), which disappeared over the course of one hour (rows 3-7). **B.** Second example neuron. As in A, this neuron responded to sTMS before cTBS. However, there was no effect of cTBS until 20 min post-stimulation (row 3). During the recovery phase, the neuron showed a period of hyperexcitability (40-50 min post-cTBS; rows 5-6).

The second example neuron also responded to sTMS before we applied cTBS (Figure 2B, top panel, notice also the weak response to light onset in the no-stimulation trials), but was not affected by cTBS in the first 10 min after cTBS (Figure 2B, second panel, sTMS-evoked response pre-compared to post-cTBS, p = 0.548). Only in the 20 and 30 min intervals after cTBS, this neuron’s excitability dropped considerably (no significant difference between stimulation and no-stimulation trials, p = 0.399 at 20 min post-cTBS; p = 0.839 after 30 min post-cTBS), although its visual response (to light onset: baseline versus post-light onset activity in no-stimulation trials; p = 2.319e-06) remained. Surprisingly, in the 40 and 50 min intervals, the excitability of this neuron temporarily increased significantly (sTMS-evoked response pre-compared to post-cTBS, p = 0.025 and p = 0.007 for the 40 and 50 min interval, respectively). Only in the 60 min interval did the neuron return to its pre-cTBS excitability level (p =0.08, pre-compared to 60 min post-cTBS in the first 40 ms after TMS onset).

A third example neuron (Figure 3A) showed yet another pattern of excitability changes after cTBS. The strong burst of activity evoked by sTMS was immediately reduced after cTBS (Figure 3A, compare top panel with second panel, p = 2.741e-04). However, this reduction in excitability grew markedly over time and reached its peak only after 50 min. At the end of our standard 60 min recording epoch, the neuron did not respond anymore to sTMS, without any sign of recovery, although we could still sporadically detect spikes (Figure 3B). In addition to the growing suppression of the sTMS response in the first hour after cTBS, we also observed a significant reduction in the baseline activity in this neuron (by 91%, p = 1.251e-05, pre-compared to 40 min post-cTBS). Because the activity of the neuron was extremely reduced, we verified that the neuron was not lost during the recording session by comparing the spike waveforms recorded before cTBS (Figure 3B upper panel) and 60 min after cTBS (Figure 3B lower panel). The firing rate of this neuron in the 60 min post-cTBS interval was reduced to barely 4 spikes/sec, but the spike waveform was virtually identical compared to the pre-cTBS epoch, which confirms that we still recorded from the same neuron. Notice also a very weak response to sTMS at 60 min post-cTBS. Overall, cTBS induced a compelling reduction in the excitability of parietal neurons, with highly variable onset time and recovery.

**Figure 3.**
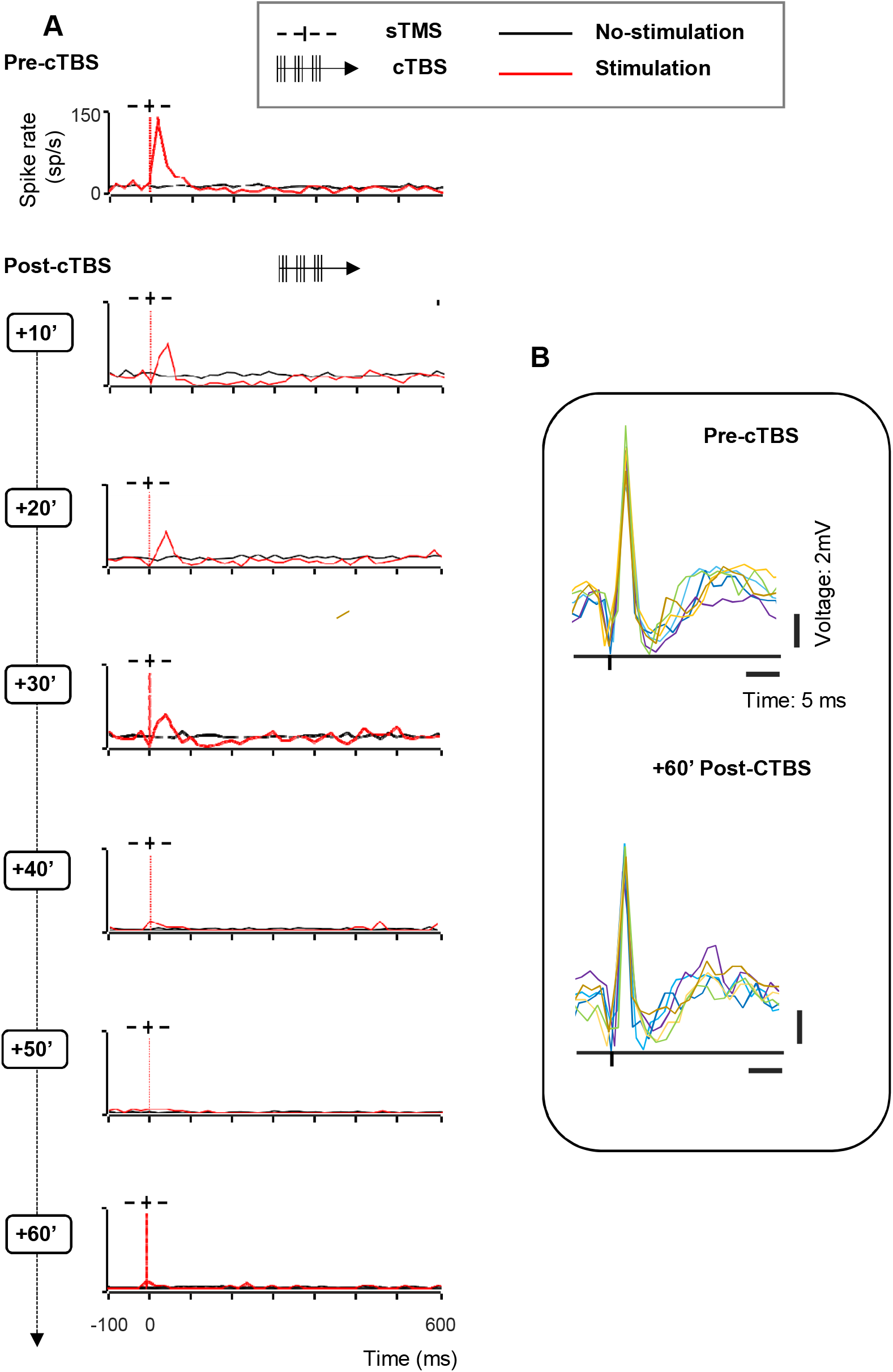
Effect of cTBS on neuronal excitability, example neuron without recovery. **A.** Spike rate of an example neuron with immediate reduced excitability after cTBS (rows 2-4). A stronger reduction, accompanied by a significant decrease of the baseline activity appeared later (40 min post-cTBS; row 4), continuing until the end of the session (60 min post-cTBS; row7). Same conventions as in Figure 2. **B.** Wave forms of the example neuron. Voltage graph showing the overlapped spike waveforms extracted from 6 consecutive trials (represented in different colors), recorded at two different time intervals (upper panel: pre-cTBS; lower panel: 60 min post-cTBS).

### Population analysis

In the first 10 minutes post-cTBS, almost half of the PFG neurons (43%; 39% in monkey Y and 49% in monkey A) showed a significant (two-sided Wilcoxon ranksum test p < 0.05) change in their sTMS response (either hypo- or hyperexcitability). However, the proportion of neurons in which cTBS affected the sTMS response gradually increased over time, such that in the 60 min post-cTBS interval, virtually all neurons (85/86) were significantly affected by cTBS (Table 1). Similarly, cTBS induced an immediate and significant change in the baseline activity (i.e. the spike rate before sTMS) in about one third of the neurons (34%, two-sided Wilcoxon ranksum test p < 0.05), which reached its maximum one hour postcTBS (93% of the neurons, Table 1). Overall, 21% of the neurons showed an effect both in the sTMS response and the baseline activity in the first 10 minutes post-cTBS, compared to 83% of the neurons showing a combined effect one hour post-cTBS.

**Table 1:** Proportions of neurons with a cTBS effect in different time epochs

|  | 10 min postcTBS | 20 min postcTBS | 40 min postcTBS | 60 min postcTBS |
| --- | --- | --- | --- | --- |
| sTMS effect | 43% (37/86)<br>Y: 20/51; A: 17/35 | 53% (46/86)<br>Y: 23/51; A: 23/35 | 86% (74/86)<br>Y: 43/51; A: 31/35 | 99% (85/86)<br>Y: 50/51; A: 35/35 |
| Baseline effect | 34% (29/86)<br>Y: 17/51; A: 12/35 | 50% (43/86)<br>Y: 24/51; A: 19/35 | 77% (66/86)<br>Y: 40/51; A: 26/35 | 93% (80/86)<br>Y: 48/51; A: 32/35 |
| Hyperexcitability in sTMS response | 10% (9/86)<br>Y: 6/51; A: 3/35 | 10% (9/86)<br>Y: 6/51; A: 3/35 | 19% (16/86)<br>Y: 10/51; A: 6/35 | 28% (24/86)<br>Y: 15/51; A: 9/35 |
| Hyperexcitability in baseline activity | 10% (9/86)<br>Y: 5/51; A: 4/35 | 12% (10/86)<br>Y: 5/51; A: 5/35 | 15% (13/86)<br>Y: 8/51; A: 5/35 | 24% (21/86)<br>Y: 13/51; A: 8/35 |

On average, cTBS significantly reduced the neuronal excitability assessed with sTMS, an effect that grew over the standard one hour post-cTBS period (Figure 4). In the first 10 min interval post-cTBS, the average sTMS response was 18% lower than pre-cTBS (p = 2.247e-08), and this reduction peaked and plateaued 40-60 min post-cTBS (24% reduction). Similarly, the reduction in the average baseline activity induced by cTBS emerged already in the first 10 min post-cTBS (by 12%, p = 6.859e-04), became more apparent 40 min post-cTBS (23 % reduction), and recovered partially at 60 min post--cTBS (18 % reduction, ANOVA on the baseline firing rate in every epoch, p = 9.604e-18, post-hoc tests 10 min compared to pre-cTBS epoch: p = 0,001, 40 min compared to pre-CTBS epoch: p = 3.706e-08, 60 min compared to pre-cTBS epoch: p = 5.101e-07). The cTBS effect was highly similar in both animals (maximum 28% reduction in sTMS response in monkey Y, and 25% reduction in sTMS response in monkey A), but the timing of the effect differed slightly, since the maximum reduction appeared at 30 min post cTBS in monkey Y and at 50 min post cTBS in monkey A (Supplementary Figure 1).

**Figure 4.**
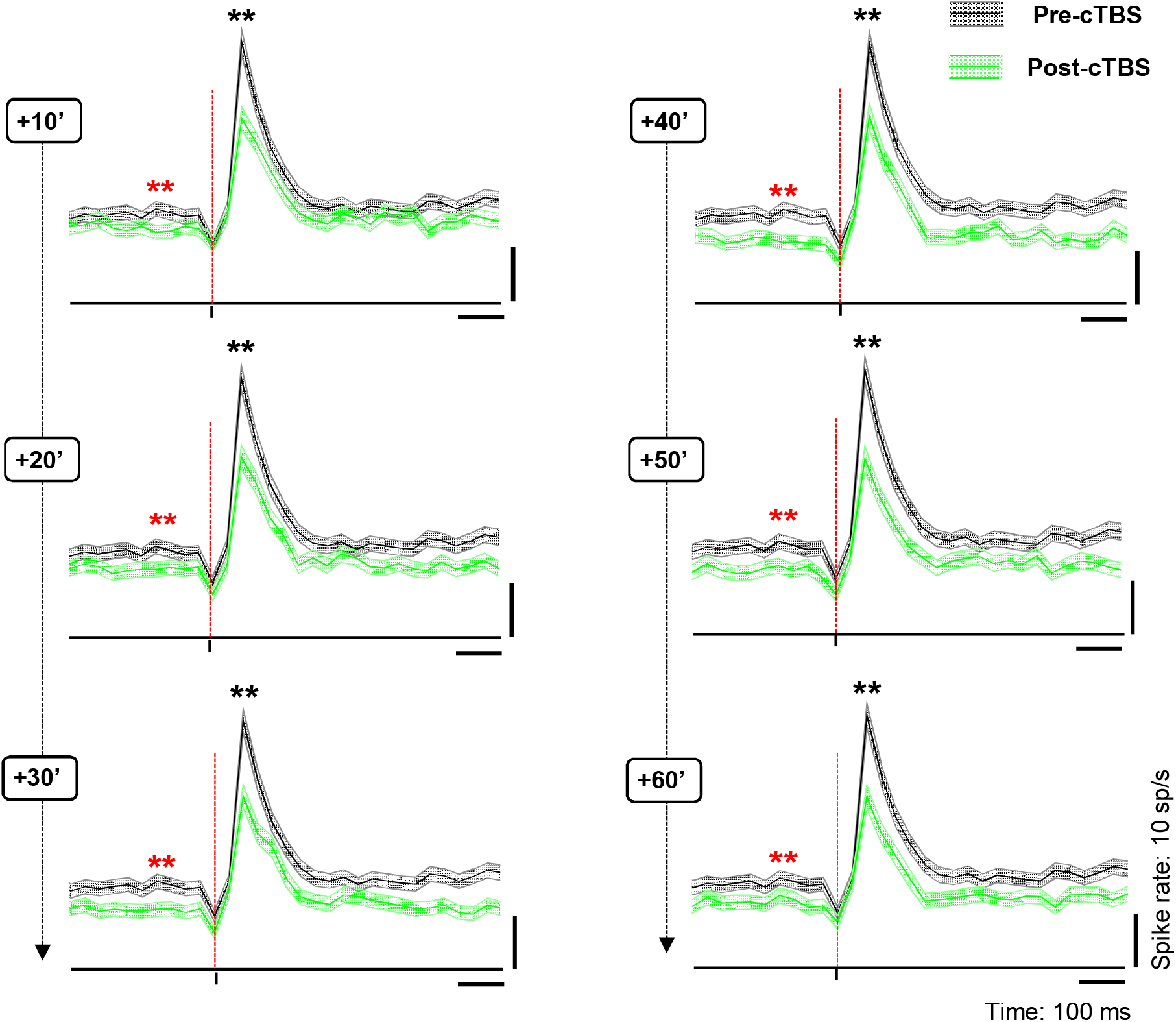
Population response to cTBS. Average sTMS responses for all neurons at different time periods post-cTBS (green) compared to pre-cTBS (black), when sTMS was applied at light onset during passive fixation. Shading displayslll ±lll the standard error. The red, dashed line indicates the sTMS pulse, aligned to light onset. The asterisks specify the statistical significance (two-sided Wilcoxon ranksum test; *p⍰≤⍰0.05; **p⍰≤⍰0.01) for changes in both the baseline activity (red) and the sTMS-evoked response (black).

To investigate whether the cTBS effect on the neuronal sTMS response and the cTBS effect on the baseline activity were based on similar neuronal mechanisms, we plotted the changes in the sTMS response and in the baseline activity against each other and calculated the correlation between the two effects in every epoch post-cTBS (Figure 5). In each individual animal (data not shown) and across all neurons combined, we measured significant and high correlation coefficients ranging from 0.54 (at 60 min post-cTBS) to 0.80 (at 30 min post-cTBS), suggesting that cTBS induced a general change in neuronal excitability, which was manifested in both sTMS response changes and alterations in spontaneous activity.

**Figure 5.**
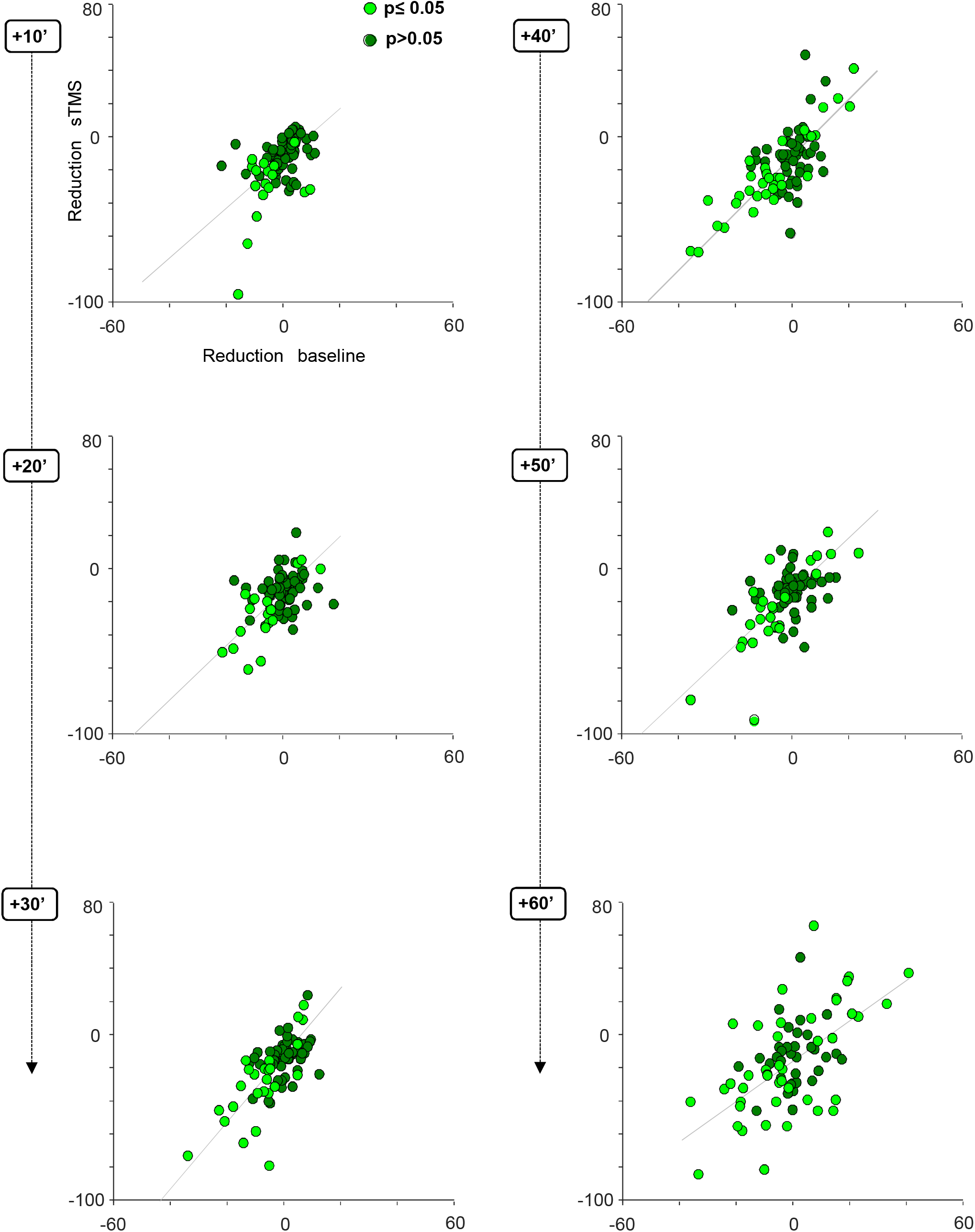
cTBS effect on neuronal excitability; scatter plots. Average response difference between the pre- and post-cTBS period plotted against the difference in the baseline activity pre-versus post-cTBS, at six consecutive, 10 min intervals post-cTBS. Each gray line indicates the least squares line (best fit). For every panel, each colored dot represents a PFG neuron. Light green dots reflect neurons showing a statistically significant (two-sided Wilcoxon ranksum test p ≤ 0.05) change in both their sTMS and baseline response (either hypo- or hyperexcitability) post-cTBS. Dark green dots indicate neurons without statistically significant effect (two-sided Wilcoxon ranksum test p > 0.05).

Figure 5 also illustrates that, although cTBS in general reduced neuronal excitability (53% of the neurons showed hypoexcitation in all time epochs post-cTBS), transient phases of hyperexcitability were not uncommon in our neuronal population. Overall, almost half of the neurons (47%) was hyperexcitable in at least one interval post-cTBS (Table 1). The large majority of these neurons (91%) were initially less excitable followed by an epoch of hyperexcitability, as the example neuron in Figure 2B. However, in a small number of neurons, hyperexcitability appeared immediately after cTBS, either as the only effect throughout the entire recording session (6%) or followed by hypoexcitability after a variable time interval (10 to 40 min, 3%). Significant increases in baseline activity were also not uncommon, since 44% of the neurons showed hyperexcitability in at least one epoch post-cTBS (Table 1).

cTBS may not only influence the average neuronal response to sTMS, but also the spike timing, i.e. the variability of the interspike intervals. We could not detect any significant effect of cTBS on the variance to mean ratio (the Fano factor) of the neurons, nor on the distribution of the interspike intervals (ISIs, data not shown). However, the power spectrum of the spike trains changed significantly after cTBS, both in the stimulation and in no-stimulation trials (Figure 6). In every epoch post-cTBS, the power in the low frequencies (below 5 Hz) was significantly reduced compared to pre-cTBS. Thus, cTBS also induced changes in the low-frequency oscillatory activity of parietal neurons.

**Figure 6.**
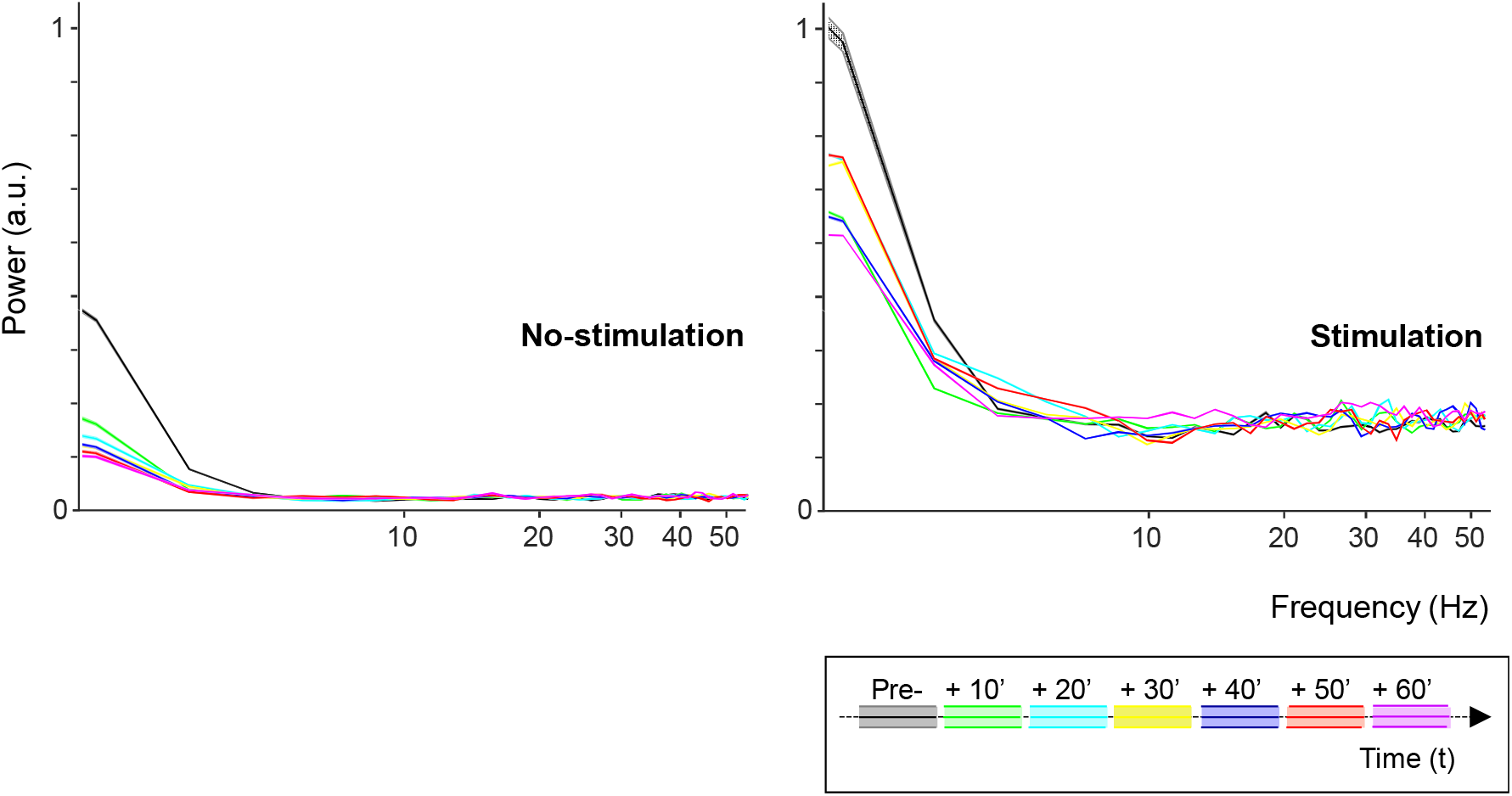
Spike oscillations analysis. Spectral power of the single-unit activity in both the no-stimulation (left panel) and stimulation (sTMS) trials (right panel), divided in 10 min intervals pre- and post-cTBS. Each color line indicates a different time interval. The shading in the graph represents ± the standard error.

### Effect of cTBS on task-related neural activity

Since a motor task may cause small movements of the electrode and since we wanted to give priority to the stability of the neural recordings, we chose to use a passive fixation task for the recordings, in which the monkeys were required to simply fixate an object illuminated in front of them to obtain a fluid reward. Despite the absence of a grasping movement, a subset of the neurons we recorded in PFG showed significant task-related activity (i.e. object responses) after light onset (N=25). The presence of object responses allowed us to test the effect of cTBS on neuronal excitability without applying sTMS. To capture all task-related responses, we tested the effect of cTBS in two intervals, an early (20 – 100 ms after light onset) and a late interval (120-520 ms after light onset). In the first 10 min epoch post-cTBS, task-related activity did not differ significantly compared to the pre-cTBS epoch (Figure 7A left panel), but in all subsequent epochs up to 60 min post-cTBS, the later task-related activity (in the interval 120-520 ms after light onset) was significantly reduced compared to pre-cTBS (Figure 7A right panel). Moreover, the early interval after light onset also showed a significant difference in the epoch 30-60 min post-cTBS. When probed with sTMS, this subpopulation of task-related neurons behaved very similar to the rest of the population, since the neuronal excitability was already reduced 10 min post-cTBS and continued to be reduced up to 60 min post-cTBS (Figure 7B). Overall, cTBS also induced a reduction in task-related activity in parietal neurons, comparable to the effects observed with sTMS.

**Figure 7.**
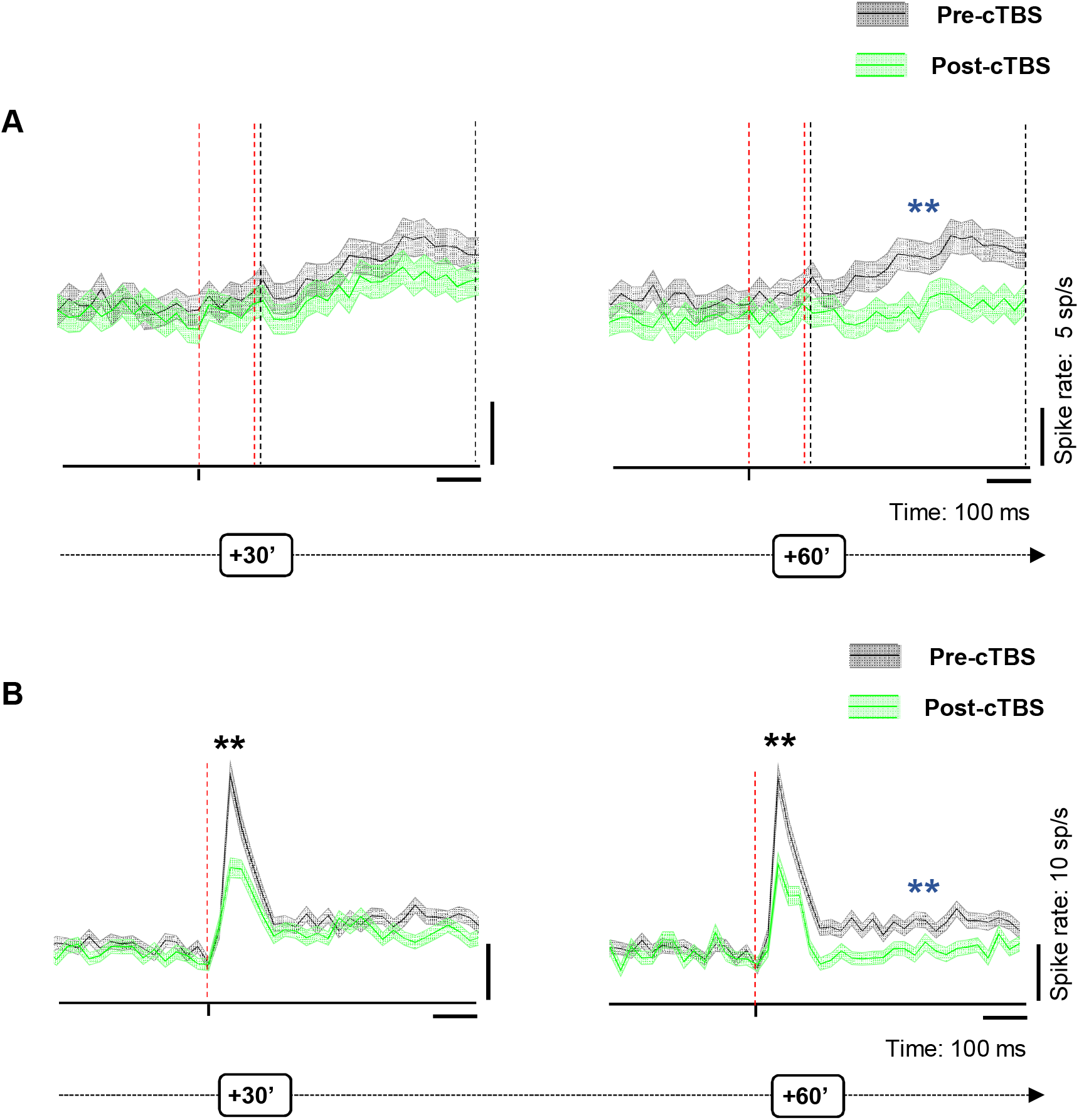
Effect of cTBS on task-related activity. **A.** Pre (black)-versus post-cTBS (green) activity in no-stimulation (sTMS) trials at two different response intervals (30 and 60 min post-cTBS). Shading displays⍰±⍰the standard error. The red, dashed line indicates the onset and offset of the first interval analyzed (early interval: 20-100 ms after light onset); the black, dashed line delimits the second interval of analysis (late: 120-500 ms after light onset). The blue asterisks specify the statistical significance (two-sided Wilcoxon ranksum test; *p⍰≤⍰0.05; **p⍰≤⍰0.01). **B.** Response of the same neuronal subpopulation during sTMS trials. Shading displays⍰±⍰the standard error. The red, dashed line indicates the sTMS pulse, aligned to light onset. The asterisks specify the statistical significance (two-sided Wilcoxon ranksum test; *p⍰≤⍰0.05; **p⍰≤⍰0.01) of the changes in both the sTMS-evoked response (black) and the post-sTMS response (task-related; blue).

### Effect of cTBS on neuronal excitability beyond one hour

Our standard recording time was one hour after cTBS (plus approximately 20 min of recordings pre-cTBS). However, in a subpopulation of neurons (N =34; Figure 8A), we could test the effect of cTBS on neuronal excitability for up to 90 min post-cTBS. Because neurons can be lost in the course of such a long time interval due to small brain movements, we only included units which the signal to noise ratio was at least 5:1 for the entire duration of the recording session, and we compared the spike waveforms pre- and post-cTBS to verify that the neuron was still present (see example spike waveforms in Figure 8B). Even 90 min after cTBS, the average sTMS response was strongly reduced (p = 1.995e-15), indicating that recovery of the excitability was at least in this subpopulation of neurons negligible or absent. Moreover, we also detected a reduction in the baseline activity and a significant reduction in neuronal activity after the sTMS-evoked burst (in the interval 200-500 ms after sTMS). A smaller subset of the neurons (N=15) was monitored up to two hours post-cTBS, but again no recovery was detectable (data not shown). Thus, the cTBS-induced reduction in neuronal excitability is very prolonged and may not recover for several hours post cTBS.

**Figure 8.**
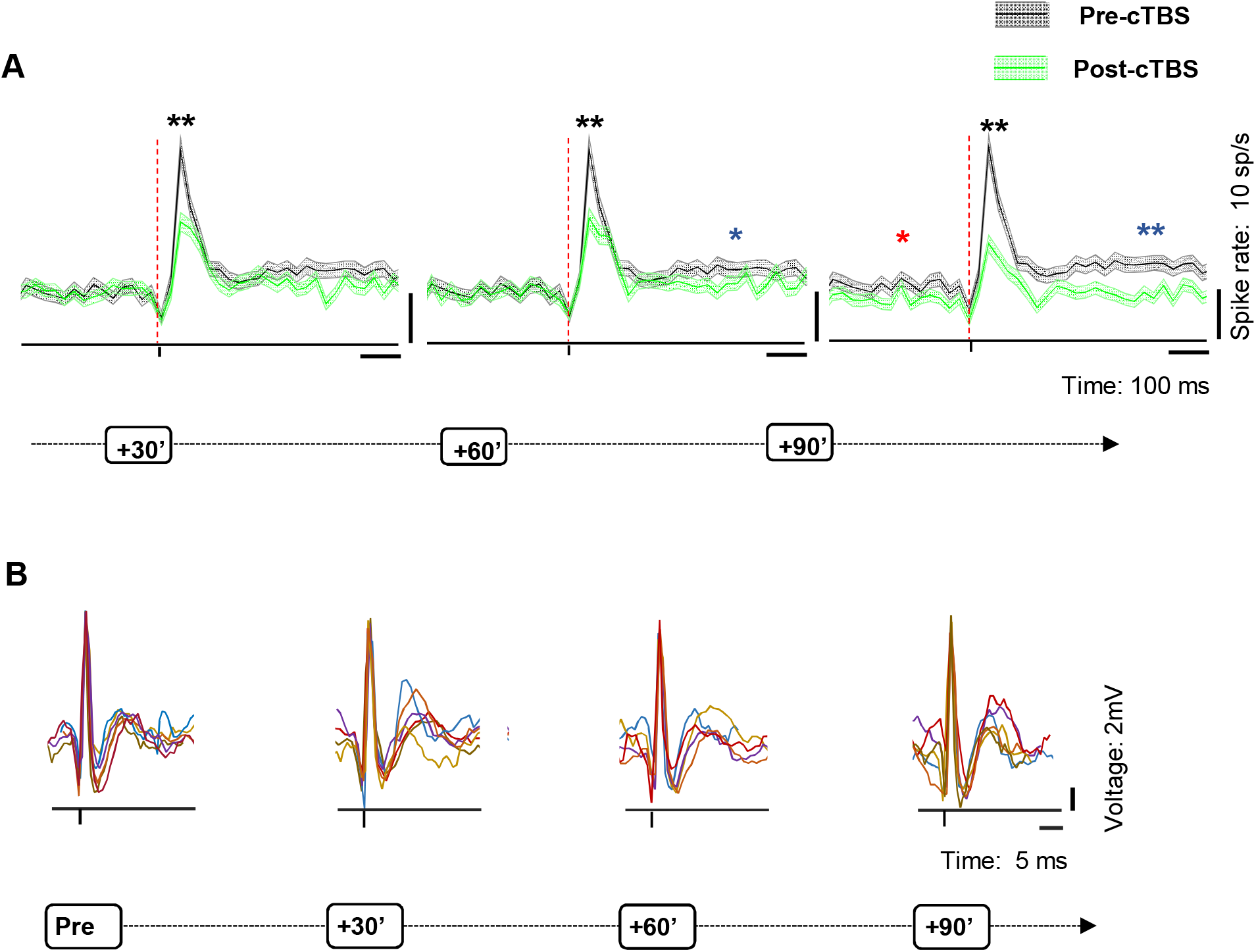
Population response to cTBS up to 90 min post-cTBS **A.** Average spike rate of all neurons (sTMS response) recorded for up to 90 min post-cTBS (green) compared to pre-cTBS (black). Shading displays⍰±⍰the standard error. Same conventions as in Figure 7. **B.** Raw signal. Example voltage graph showing the pre- and post-cTBS spike waveforms from a representative neuron, monitored over time. In each panel, we plotted 6 consecutive spike waveforms (represented in different colors), recorded at four different time intervals (from left to right: pre-cTBS and 30, 60 and 90 min post-cTBS).

### Behavioral effect of cTBS on grasping

To verify that our cTBS protocol (80% of the RMT, applied for 20 sec) causally interfered with grasping behavior and to monitor the behavioral effect of cTBS over time, we trained two other rhesus monkeys in a simple visually-guided grasping task and measured their grasping time (i.e. the time elapsed between the lift of the hand and the pull of the object) before and up to two hours after cTBS (at 80% of the RMT) over parietal cortex (Figure 9). To control for potential aspecific effects related to the mounting of the coil or the sound elicited by cTBS, we also included a low-intensity cTBS (at 40% of the RMT) condition. In both animals (Figure 9A-B), high-intensity cTBS induced a significant (compared to no-stimulation) increase in the grasping time in the first 20 min post cTBS, which grew until 40-60 min post cTBS and remained constant thereafter (repeated measures ANOVA with stimulation level and epoch as factors, interaction p =7.312e-22 for monkey P and p = 3.238e-14 for monkey D). At the peak of the effect, the grasping time increased by approximately 50 ms in monkey P (an 11% increase) and 40 ms in monkey D (12% increase). Low-intensity cTBS did not induce a consistent change in the grasping time (ANOVA, p = 1.02e-0.7). Note that, similar to the neural recordings, we did not observe any recovery of the grasping time even two hours post-cTBS. Thus, cTBS applied at 80% of the RMT for 20 sec not only induced a strong reduction in neural excitability, but also a prolonged behavioral deficit in an object grasping task.

**Figure 9.**
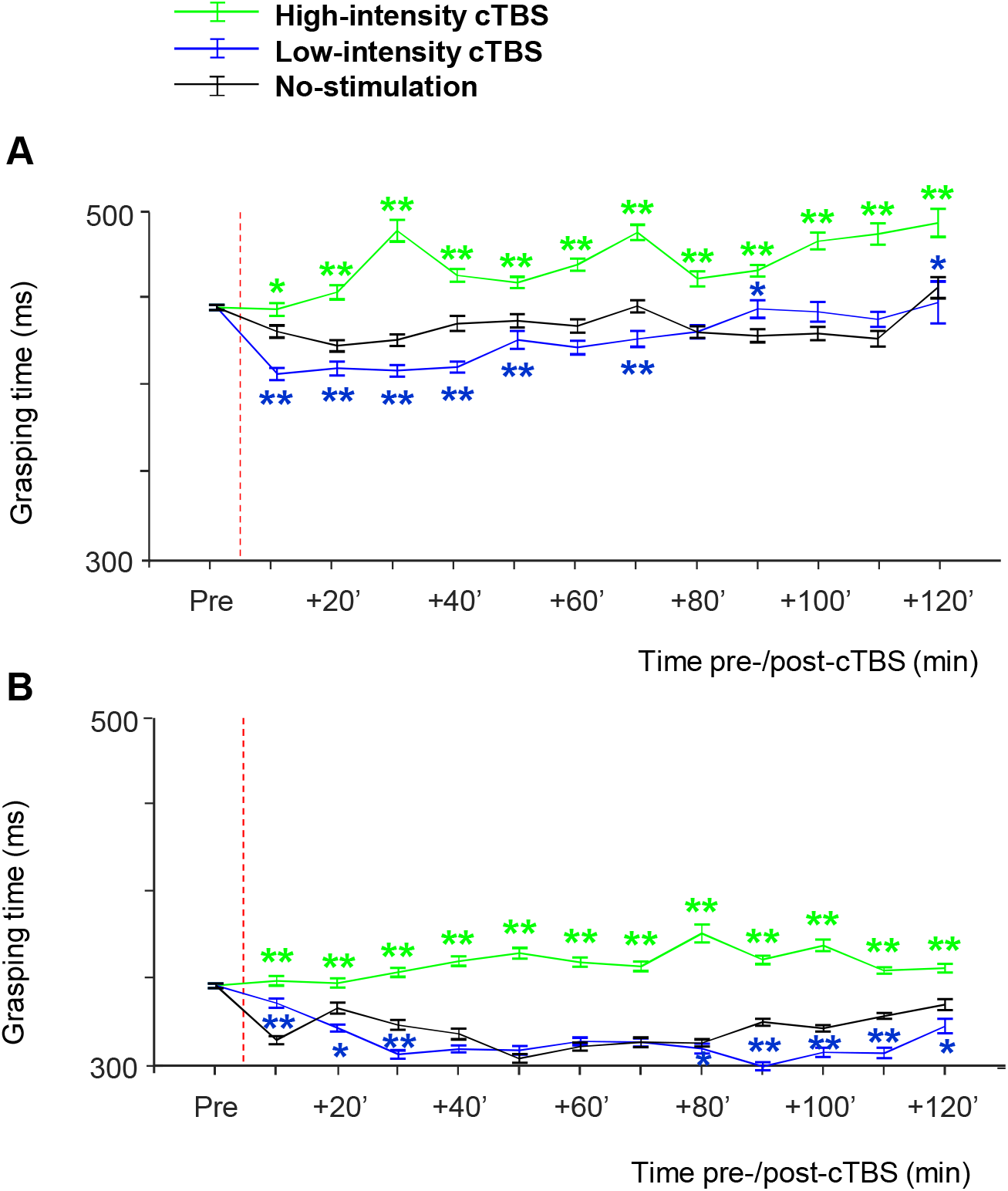
Behavioral effect of cTBS, experiment 2. **A.** Normalized grasping times for high-intensity (solid, green line), low-intensity (solid, blue line) and no-stimulation (solid, black line) cTBS as a function of time in monkey P. Grasping times of minimally 1700 correct trials were averaged in time-epochs of 10 min. The red, dashed line represents the stimulation onset (cTBS). The asterisks indicate the level of statistical significance (two-sided Wilcoxon Ranksum test, *: p < 0.05, **; p < 0.01). **B.** Same results for monkey D.

## DISCUSSION

We observed a pronounced and long-lasting reduction in neuronal excitability after cTBS in macaque parietal neurons, which was mirrored in a behavioral impairment when grasping objects. This reduced excitability of individual neurons grew over time, consistent with previous studies in humans, but some neurons exhibited periods of hyperexcitability during the recovery phase. These results provide the first experimental evidence on the neural effects of cTBS on single neurons in awake behaving monkeys.

Our basic findings, both at the single-cell level and at the behavioral level, are remarkably consistent with the known effects of cTBS over primary motor cortex in human volunteers (Huang et al., 2005). The average amplitude of the MEP – a measure of neuronal excitability in motor cortex – is unaltered in the first minutes after cTBS, then gradually declines and recovers 25 (after 20 sec cTBS) or 60 min (after 40 sec cTBS) later. In parallel, the reaction time significantly increases 10 min after cTBS over primary motor cortex (Huang et al., 2005). The time course of the neural and behavioral effects we measured after 20 sec of cTBS in monkeys seem to be more similar to the 40 sec cTBS protocol in humans, possibly due to the thinner skull of monkeys. Nevertheless, it should also be noted that we applied cTBS over parietal cortex, whereas most cTBS studies in humans have targeted primary motor cortex (but see Tunik et al., 2005). Overall, the consistency of our findings with the human literature strongly suggests that single-cell recordings after cTBS in awake macaque monkeys represent a valid approach to understand the neural effects of cTBS.

The crucial advantage of our approach is that we can measure the wide range of changes in excitability in individual neurons, while the MEP amplitude represents an excitability measure of a large population of neurons. In addition, we could demonstrate that the inhibitory influence of cTBS progressively recruits more neurons over time, even neurons that were initially unaffected. Finally, we showed that cTBS induces a general reduction in neuronal excitability, which appears in changes in baseline firing rate, sTMS-evoked responses, low-frequency oscillatory activity and task-related activity.

Extracellular recordings do not easily allow determining the cell type of the unit that is being recorded based on the spike waveform (see for example Woloszyn and Sheinberg (2012) compared to Vigneswaran et al., 2011i), nor the cortical layer in which these units were recorded. In general, we recorded from neurons immediately under the TMS coil generating large spike waveforms, that we could monitor for more than one hour and up to two hours post-cTBS. However, the large range of effects we observed in our population of neurons may suggest that cTBS exerts specific effects on different cell types, on neurons in different layers, and in a different orientation and/or location with respect to the TMS coil. In nonhuman primates, future studies could address these questions with advanced techniques such as calcium imaging (Ikezoe et al., 2013; Tang et al., 2020).

Previous studies have suggested that cTBS may induce long-term depression (LTD)-like effects on cortical synapses (Huang et al., 2011). LTD, a widespread phenomenon driving synaptic plasticity both in subcortical structures and in the cortex, is typically induced by low-frequency stimulation (LFS, at 1 Hz), and its underlying molecular mechanisms may be very diverse depending on brain area and developmental stage (Massey and Bashir, 2007). In visual cortex, 15 min of 1 Hz stimulation induces LTD of synaptic responses (Kirkwood and Bear, 1994). However, the LTD effect studied in cortical slices appears immediately after the end of LFS and remains constant for up to half an hour. In contrast, the reduction in neuronal excitability we observed after cTBS grew gradually and reached its maximum 30-50 min after cTBS, similar to the reduction in the amplitude of the MEP observed after cTBS in humans (Huang et al., 2005). Almost half of the neurons in our sample were not affected at all in the first epoch post-cTBS, but became less excitable in the hour following cTBS. Therefore, the time course of the cTBS effect we measured does not seem to be compatible with a pure LTD effect. The gradual reduction in neuronal excitability seems more consistent with an increase in the local concentration of the inhibitory neurotransmitter GABA, which may slowly spread in the cortical area under the TMS coil. It is noteworthy that magnetic resonance spectroscopy has demonstrated an increase in GABA in human motor cortex after cTBS (Stagg et al., 2009). In the nonhuman primate model, future studies will be able to investigate in more detail which molecular mechanisms are responsible for the effect of cTBS.

Unexpectedly, some neurons also showed a period of hyperexcitability after an initial phase of reduced excitability caused by cTBS. We interpret these findings as evidence that nearby inhibitory interneurons showed a different time course of recovery after cTBS, such that at some point post-cTBS the neuronal excitability had recovered while the normal inhibitory inputs to the neuron were still less active. In a previous study (Romero et al., 2019), we also observed that TMS affects both large pyramidal neurons and small (inhibitory) interneurons. The temporary hyperexcitability in our data may also be related to the rare occurrence of seizures after TMS (Lerner et al., 2019).

Because we concentrated on measuring the changes in neuronal excitability after cTBS, we did not map the area of cortex under the coil that was affected by cTBS. Single-pulse TMS affects a surprisingly small volume of cortex, which we estimated to be not larger than 2 by 2 by 2 mm (Romero et al., 2019). If our hypothesis of the spreading of GABA is correct, we expect that a slightly larger volume of cortex will be affected by cTBS. Moreover, the robustness of the behavioral deficit induced by cTBS suggests that we inactivated a larger region in parietal cortex.

The neural effects of cTBS were highly similar in the two animals we used, whereas the effects of cTBS in human volunteers are notoriously variable (Hamada et al., 2013; Hordacre et al., 2017; Jannati et al., 2017). The reproducibility of our results was most likely related to the very controlled conditions in which we applied cTBS. Most importantly, the TMS coil was rigidly anchored to the head implant of the animal, so that we kept both the position and the orientation of the coil similar across sessions.

The critical advantage of the macaque monkey model is that we can measure single-cell activity and behavioral performance before and after cTBS. In a subset of neurons that we could record for more than one hour, we did not observe any recovery of the excitability even after two hours. Consistent with this observation, the behavioral effect of cTBS also lasted at least two hours. Therefore, our cTBS paradigm in monkeys may have caused longer-lasting effects than in human volunteers, possibly due to the thinner skull of the monkey. We did not observe any effect on the neuronal excitability on consecutive days, since we could always record single-unit activity over several weeks of recordings.

Investigating the neural effects of non-invasive brain stimulation techniques requires adequate animal models, so that behavioral measurements and detailed recordings of individual neurons can be combined with neuromodulation. In future studies, other stimulation protocols such as intermittent theta-burst stimulation and other neuromodulation techniques such as transcranial Alternating Current Stimulation can be investigated and optimized in animal models using a similar approach.

---

**Supplementary Figure 1.**
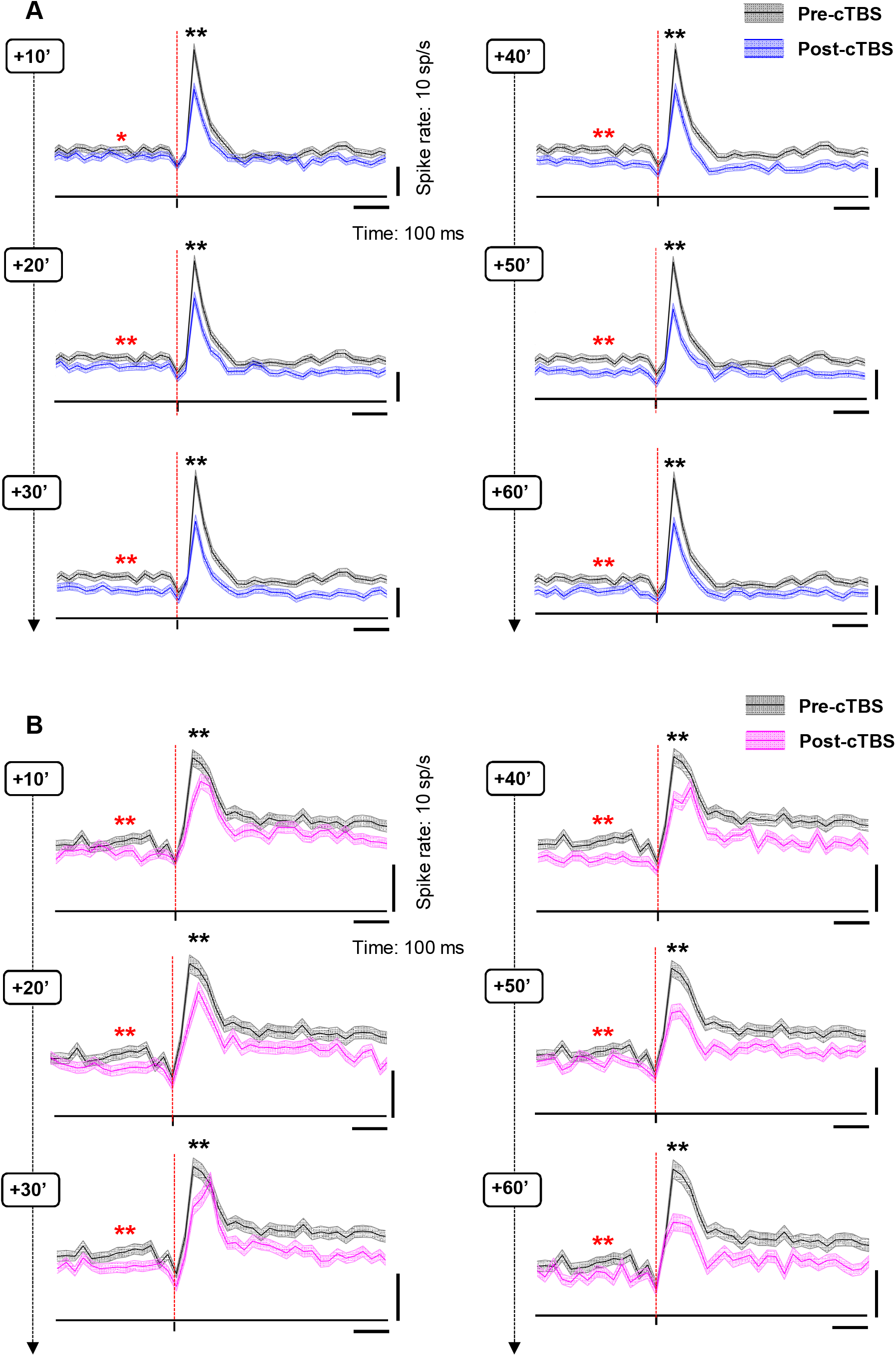
Population response to cTBS for monkeys Y and A, separately. **A.** Average spike rate of the sTMS response for all neurons recorded in monkey Y at different time periods post-cTBS (blue) compared to pre-cTBS (black), when sTMS was applied at light onset during passive fixation. Shading displays⍰±⍰the standard error. The red, dashed line indicates the sTMS pulse, aligned to light onset. The asterisks specify the statistical strength (two-sided Wilcoxon ranksum test; *p⍰≤⍰0.05; **p⍰≤⍰0.01) for changes in both the baseline activity (red) and the sTMS-evoked response (black). B. Equivalent graph for monkey A, plotting the average spike rate of the sTMS response recorded at different time periods post-cTBS (magenta) compared to pre-cTBS (black). Same conventions as in A.

## ACKNOWLEDGEMENTS

This work was supported by Fonds voor Wetenschappelijk Onderzoek Vlaanderen (Odysseus grants G.0007.12 and G.0C51.13N), and Program Financing (PFV10/008). We would like to thank Stijn Verstraeten, Christophe Ulens, Piet Kayenbergh, Gerrit Meulemans, Marc De Paep, Astrid Hermans and Inez Puttemans for their technical contributions.

